# No Miracle, Just a Mineral: The Not-So-Magical Antimicrobial World of Chlorine Dioxide

**DOI:** 10.1101/2024.06.03.597186

**Authors:** R. Dudek-Wicher, M. Brożyna, J. Paleczny, B. Mączyńska, B. Dudek, P. Migdał, A. Dołowacka-Jóźwiak, J. Fischer, A. Junka

## Abstract

This study evaluates the *in vitro* antimicrobial efficacy and cytotoxicity of acidified sodium chlorite (ASC), a source of chlorine dioxide. Despite its controversial promotion in alternative medicine as a cure-all solution, known as “Miracle Mineral Solution” (MMS), the data on its factual medicinal activity is very limited. Therefore, we aimed to elucidate the activity of ASC against biofilms of *Staphylococcus aureus, Pseudomonas aeruginosa, Enterococcus faecalis, Streptococcus mutans, Pseudomonas aeruginosa, Escherichia coli*, and *Lactobacillus sp*. The study also investigated the i*n vitro* cytotoxic effects of ASC towards eukaryotic fibroblasts and in *in vivo* model of *Galleria mellonella* larvae. Our findings demonstrate that these concentrations of ASC which can effectively eradicate biofilms, also pose potential health risks due to their *in vitro* and *in vivo* cytotoxicity. It implies that ASC applied in human can lead to damage to the mucous membrane of the gastrointestinal tract. This research contributes to the ongoing debate on the safety and efficacy of chlorine dioxide in clinical applications, highlighting the need for precise dosing to avoid mucosal damage in therapeutic contexts.

## 1. Introduction

Sodium chlorite is an inorganic compound utilized in plethora of applications [1]. Acidification of sodium chlorite neutral solutions results in the formation of acidified sodium chlorite (ASC) and the stabilization of the pH value. During the process, a significant quantity of chlorine dioxide (ClO_2_) is liberated:

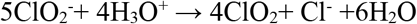

Sodium chlorite solutions are frequently marketed in alternative medicine circles as ‘Miracle Mineral Solution’ (MMS); a panacea for a wide array of illnesses. Distributors of MMS advocate that when mixed with citric acid, it exhibits efficacy against autism and cancer, as well as against various microbial pathogens including COVID-19 [2-4].

The ClO_2_ is also a potent bleaching agent that is not intended for ingestion, because of potential to cause significant cellular and tissue damage and life-threatening adverse reactions [5-11]. Despite these warnings, the MMS product remains available for purchase through various online retailers [12-16]. From the other hand, aqueous solutions of ASC may be applied as antimicrobial agent but only under strictly regulated concentrations (e.g. 0.003% to 0.016% concentrations (wt/vol %) ClO_2_) and conditions, as evidenced by numerous clinical studies and approvals from regulatory bodies [17-24].

Therefore, our study aims to identify the effective antimicrobial/antibiofilm concentrations of ASC and apply it in the next step towards *in vitro* cell lines/ *in vivo* model to assess the potential cytotoxic effects. The superior aim of this research is to provide empirical, scientific data on the efficacy and safety of ASC. This is crucial in either validating or refuting the claims made in alternative medicine circles. If the research finds specific concentrations and conditions under which ASC is effective and safe, it could lead to its more controlled and informed use in medical settings and managing infections. Conversely, if the results show limited efficacy or significant toxicity, it could help debunk unwarranted claims and protect patients and consumers from potentially harmful health advice.

## 2. Materials and methods

The following 7 solutions were applied:

A. Hydrochloric acid (HA), (5%, Chempur, Poland)
B. Gluconic acid (GA), (50%, Chempur, Poland)
C. Sodium chlorite (SC), (25%, Chempur, Poland)
D. Prontosan – 0,1% polyhexamethylene biguanide (PHMB, B.Braun, Germany)
E. ASC1 – acidified sodium chlorite solution acidified with HA
F. ASC2 – acidified sodium chlorite solution acidified with GA

SC (25%) was diluted in a ratio of 1:1 with 5% HA (ASC1) or 50% GA (ASC2). After 60s of activation, deionized water was added to interrupt the hydrolysis process. The final ASC1 working solution contained 0.53% SC and 0.16% HA (pH=2.5). ASC2 contained 0.53% SC and 1.6% GA (pH=3.12). The concentration of ClO_2_ in ASC1 and ASC2 was 0.383% (3830ppm). Control solutions of 0.53% SC (pH=7.86); 0.16% HA (pH=1.63); 1.6% GA were prepared (pH=2.86). pH measurement was performed with Pehametr Elmetron CPI-505 (Emetron, Poland).

The research was performed on reference strains (American Type Culture Collection (ATCC, Manassas, VA, USA; and Polish Collection of Microorganisms (PCM, Poland)):

- *Staphylococcus aureus* ATCC 6538 - (MSSA - Methicillin sensitive)
- *Staphylococcus aureus* ATCC 33591 - (MRSA - Methicillin resistant)
- *Enterococcus faecalis* ATCC 25212
- *Lactobacillus rhamnosus* PCM 489
- *Lactobacillus casei* PCM 2639
- *Pseudomonas aeruginosa* ATCC 15442
- *Escherichia coli* ATCC 25922
- *Streptococcus mutans* ATCC 25175

### 2.1. Evaluation of the Minimal Inhibitory Concentration (MIC)

The strains were transferred from agar plates to appropriate liquid broths and incubated at 37°C/5%CO_2_/24h. The standard 96-well plate microdilution method with use of tetrazole chloride was performed. The PHMB served as positive control (C+). Growth and sterility control and control tests with HA, GA, and SC were also performed. The highest tested concentrations were HA: 0.08%; GA: 0.08%; SC: 0.27%; PHMB: 0.05%. All procedures were performed in triplicates. The microbial turbidity studies were performed using Thermo Scientific Multiskan Go spectrometer (Thermo-Fischer Scientific, Finland) with a wavelength of 600nm. MIC values were expressed as the percentage (%ClO_2_) and part per million (ppm) concentration of ClO_2_.

### 2.2. Visualization with Transmission electron microscopy (TEM)

The samples of MSSA and *P. aeruginosa* were fixed in glutaraldehyde (POCH, 2.5%) and subjected to centrifugation (5min, 50µf). The samples were then passed through an ascending alcohol series and embedded in a medium-hard epoxy resin. After polymerization, ultra-thin sections were prepared on an ultramicrotome (Leica). Sections of 60nm were prepared from the resin blocks and placed on copper grids (400 Mesh) with formvar film and carbon coating. The contrasting was performed using 2% uranyl acetate (MicroShop, Poland) (10 min) and 2% osmium tetroxide (Agar Scientific, UK) (2h) as described elsewhere [25]. Imaging was performed using a JEOL 1200, (JEOL, Japan) transmission microscope.

### 2.3. Evaluation of the Minimal Biofilm Eradication Concentration (MBEC)

Suspensions of density 10^5^ of each species were prepared as described in 2.1 section. 200mL of each suspension was added to the wells of 96-well plate and incubated at 37°C/5%CO_2_/24h. Next, a series of tested compounds’ dilutions in appropriate liquid media were prepared. The highest tested ClO_2_ concentration was 0.383% (3830ppm) obtained from both formulations: ASC1 and ASC2. The highest tested concentrations of control substances were HA: 0.16%; GA: 0.16%; SC: 0.53%; PHMB: 0.1%. The analyses were performed as we described earlier [26]. The absorbance was measured with a wavelength of 490nm. MBEC values were expressed as %ClO_2_ and ppm of ClO_2_. The tests were performed in triplicates.

### 2.4. Visualization of biofilm with fluorescence microscopy

Concentrations equal MBEC, <MBEC, and >MBEC of ASC1 were applied against tested biofilms preformed in the wells of 24-well plates. To visualize live cells, positive control of each chosen strain was prepared. Plates were incubated at 37°C/5%CO_2_/24h. Then biofilms were dyed with the mixture of SYTO-9 and propidium iodide (PI) dyes (FilmTracer™ LIVE/DEAD® Biofilm Viability kit, Invitrogen, Ltd., Paisley PA4 9RF, UK) to visualize live and dead cells, respectively. The samples were visualized using Lumascope 620 (Etaluma, Carlsband, CA, USA) with 20µm magnification objective Olympus IPC phase (Shinjuku, Japan). The field of vision recorded was 0.49mm and the frame size was 1200 x 1200 pixels. The excitation/emission wavelengths for SYTO9 and propidium iodide were 480/500nm and 490/635nm, respectively.

### 2.5. Biofilm Eradication from Hydroxyapatite (HAP) Surface

MBEC concentrations of ASC1 were applied against 24-hour old biofilms preformed on hydroxyapatite discs (HAPd) (manufactured as described by Junka et al.) in the wells of a 24-well plate [27] in 37°C/5%CO_2_/24h conditions. Then, all samples were stained, and the analyses were performed as we described earlier [26]. The absorbance was measured with a wavelength of 490nm.

### 2.6. Visualization of biofilm with Scanning Electron Microscopy (SEM)

MSSA, *P. aeruginosa, S. mutans* and *L. rhamnosus* biofilms formed on HAPd were selected for this stage of research. Biofilms were fixed in 2.5% glutamate aldehyde (Carl Roth, Germany) for 24h/4 °C. Then, the samples were rinsed two times with a 0.2M cacodyl buffer to remove any residuals. The dehydration process was conducted with increasing concentrations of ethanol (30, 60, 80, 90, and 9.99%). The obtained samples were coated with a 15nm layer of carbon using a high vacuum carbon coater (ACE 600, Germany) and imaged with the ZEISS Auriga 60 scanning electron microscope (Zeiss, Germany).

### 2.7. Evaluation of antibiofilm activity of ASC after different activation time

SC (25%) was diluted in a ratio of 1:1 with 5% HA or 50% GA. At this stage different acidification times were applied: 60s, 120s, 360s before distilled, deionized water was added to tested mixtures. Next, all steps of this stage of the study were performed as described in section 2.3.

### 2.8. Cytotoxicity of Tested Solutions

#### A. *In vitro* - towards L929 Fibroblast Cells

The standard neutral red cytotoxicity test was performed. A 100mL of suspension of fibroblasts L929 (American Type Culture Collection, Rockville, MD, USA) of 10^5^ cells/mL density was added to 96-well plate and incubated for 24h/37°C. Next, the medium was removed and MIC and MBEC concentrations of ASC1 and ASC2 were poured (100µL/well) [28].

#### B. *In vivo* - in model of *Galleria mellonella* larvae

The *in vivo* model of *Galleria mellonella* larvae was applied to perform analysis of cytotoxicity of ASC1 in A-0.383% and B- 0.002992% concentrations. The control setting was the PHMB, 75% ethanol (ChemPur, Poland) and Dulbecco’s Phosphate Buffered Saline (PBS; Biowest, Riverside, MO, USA). The *G. mellonella* larvae in a stadium of sixth instar (weight of 0.21g), were selected. 20µL of compounds mentioned above were injected into the larvae. Each compound was injected totally into 20 larvae in two separate experiments (2x10 larvae). The larvae were then incubated, and their mortality was monitored after 2h from injection and subsequently every 24 hours up to 120th hour. Death was stated when the larvae were nonmobile, melanized, and did not react to physical stimuli.

## 3. Results

### 3.1. Minimal Inhibitory Concentration evaluation (MIC)

The MICs of ASC1 and ASC2 towards *P. aeruginosa* were lower than the MIC of control antiseptic agent - PHMB. For *E. coli*, the opposite results were obtained. The lowest MIC of ASC1 was observed towards MSSA while the lowest MIC of ASC2 was revealed for MSSA but also for *E. coli*. S. *mutans, L. rhamnosus* and *L. casei* were the most tolerant to ASC1 than to ASC2. ASC2 has shown greater antimicrobial activity against *E. coli* and *L. casei* than ASC1. The antimicrobial activity of ASC2 against *L. casei* was like the activity of PHMB. In the case of *S. mutans, E. faecalis* and *L. casei*, antimicrobial activity of ASC1 and ASC2 did not exceed the activity of SC (**Table 1**).

**Table 1.** The MIC (%; ppm) of tested solutions

| Tested strain/Tested solution | MSSA |  | MRSA |  | *E. faecalis* |  | *S. mutans* |  |
| --- | --- | --- | --- | --- | --- | --- | --- | --- |
|  | % | ppm | % | ppm | % | ppm | % | ppm |
| **ASC1**<br>(%ClO<sub>2</sub>) | 0.002992 | 29.92 | 0.005984 | 59.84 | 0.005984 | 59.84 | 0.023938 | 239.38 |
| **ASC2**<br>(%ClO<sub>2</sub>) | 0.002992 | 29.92 | 0.005984 | 59.84 | 0.005984 | 59.84 | 0.023938 | 239.38 |
| **HA** | 0.08 | 800 | 0.08 | 800 | 0.08 | 800 | 0.08 | 800 |
| **GA** | 0.8 | 8000 | 0.8 | 8000 | 0.8 | 8000 | 0.8 | 8000 |
| **SC** | 0.0083 | 83 | 0.0083 | 83 | 0.0041 | 41 | 0.0165 | 165 |
| **PHMB (C+)** | 0.000098 | 0.98 | 0.000098 | 0.98 | 0.00039 | 3.9 | 0.000098 | 0.98 |
| Tested strain/Tested solution | *P. aeruginosa* |  | *E. coli* |  | *L. rhamnosus* |  | *L. casei* |  |
|  | % | ppm | % | ppm | % | ppm | % | ppm |
| **ASC1**<br>(%ClO<sub>2</sub>) | 0.005984 | 59.84 | 0.005984 | 59.84 | 0.023938 | 239.38 | 0.023938 | 239.38 |
| **ASC2**<br>(%ClO<sub>2</sub>) | 0.005984 | 59.84 | 0.002992 | 29.92 | 0.011969 | 119.69 | 0.005984 | 59.84 |
| **HA** | 0.08 | 800 | 0.08 | 800 | 0.08 | 800 | 0.08 | 800 |
| **GA** | 0.8 | 8000 | 0.8 | 8000 | 0.8 | 8000 | 0.8 | 8000 |
| **SC** | 0.0083 | 83 | 0.0041 | 41 | 0.0165 | 165 | 0.0165 | 165 |
| **PHMB (C+)** | 0.00625 | 62.5 | 0.000195 | 1.95 | 0.00625 | 62.5 | 0.00625 | 62.5 |

### 3.2. Visualization of cells treated with MIC concentration with TEM

The TEM analysis performed on Gram-positive *S. aureus* and Gram-negative *P. aeruginosa* confirmed that ASC1 applied in MIC concentration mechanism of action relies on the destruction of cell walls/membranes leading to the cells’ deformation, cytoplasm leakage, and to death of cells (**Figure 1**).

**Figure 1.**
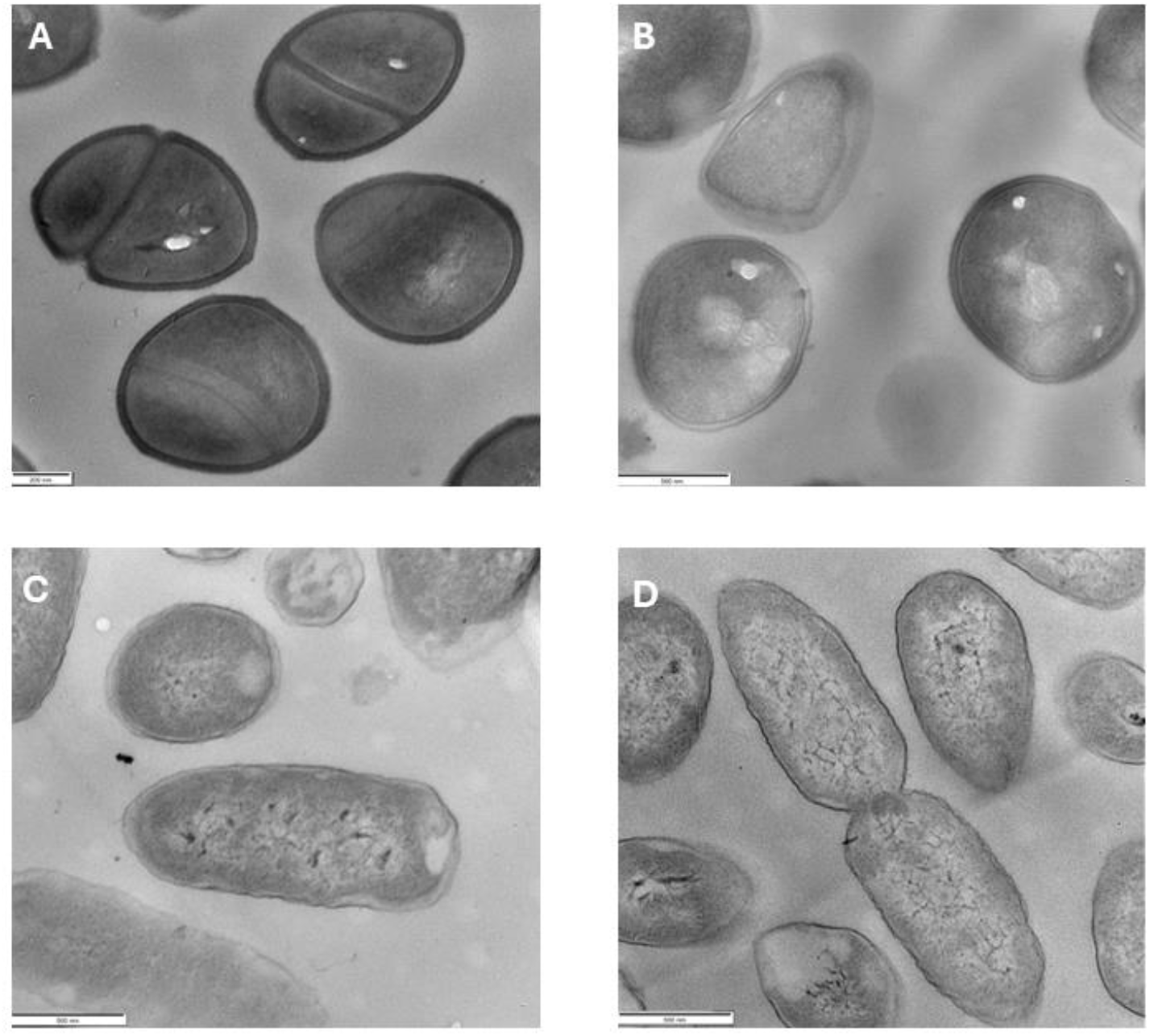
Visualization of *S. aureus* (A, B) and *P. aeruginosa* (C, D) treated (B, D) or non-treated (A, C) with MIC solution of tested solution. The deformation of cellular shapes and cell wall as well as loss of density of microbial cytoplasm (areas of lower electron density visualized by light grey color) are explicitly visible in the images of treated cells **(C, D)**.

### 3.3. Minimal Biofilm Eradication Concentration evaluation (MBEC)

None of the tested solutions were more effective against tested biofilms than PHMB. SC has shown greater antibiofilm activity against all species than ASC1 and ASC2. In turn, ASC2 displayed greater antibiofilm potential than ASC1. Biofilms formed by P. *aeruginosa* and *L. casei* were the most resistant to both ASC1 and ASC2 (**Table 2**).

**Table 2.** The MBEC of tested compounds (%; ppm)

| Tested strain/Tested solution | MSSA |  | MRSA |  | *E. faecalis* |  | *S. mutans* |  |
| --- | --- | --- | --- | --- | --- | --- | --- | --- |
|  | % | ppm | % | ppm | % | ppm | % | ppm |
| ASC1 (%ClO<sub>2</sub>) | 0.1915 | 1915 | 0.1915 | 1915 | 0.1915 | 1915 | 0.09575 | 957.5 |
| ASC2 (%ClO<sub>2</sub>) | 0.047875 | 478.75 | 0.09575 | 957.5 | 0.09575 | 957.5 | 0.047875 | 478.75 |
| HA | 0.16 | 1600 | 0.16 | 1600 | 0.16 | 1600 | 0.08 | 800 |
| GA | 1.6 | 16000 | 1.6 | 16000 | 1.6 | 16000 | 0.4 | 4000 |
| SC | 0.033125 | 331.25 | 0.06625 | 662.5 | 0.0165 | 165 | 0.0083 | 83 |
| PHMB (C+) | 0.0125 | 125 | 0.00625 | 62.5 | 0.003125 | 31.25 | 0.001563 | 15.63 |

| Tested strain/Tested solution | *P. aeruginosa* |  | *E. coli* |  | *L. rhamnosus* |  | *L. casei* |  |
| --- | --- | --- | --- | --- | --- | --- | --- | --- |
|  | % | ppm | % | ppm | % | ppm | % | ppm |
| ASC1 (%ClO<sub>2</sub>) | 0.383 | 3830 | 0.09575 | 957.5 | 0.1915 | 1915 | 0.383 | 3830 |
| ASC2 (%ClO<sub>2</sub>) | 0.383 | 3830 | 0.09575 | 957.5 | 0.09575 | 957.5 | 0.383 | 3830 |
| HA | 0.16 | 1600 | 0.16 | 1600 | 0.16 | 1600 | 0.08 | 800 |
| GA | 1.6 | 16000 | 1.6 | 16000 | 1.6 | 16000 | 0.8 | 8000 |
| SC | 0.033125 | 331.25 | 0.033125 | 331.25 | 0.0165 | 165 | 0.033125 | 331.25 |
| PHMB (C+) | 0.00625 | 62.5 | 0.00625 | 62.5 | 0.003125 | 31.25 | 0.00625 | 62.5 |

### 3.4. Visualization of biofilm with fluorescence microscopy

After applying the MBEC concentrations, the quantity of live cells in biofilms significantly decreased compared to positive control (**Figure 2**). *P. aeruginosa* biofilm was one of the most resistant for ASC1 treatment which is consistent with MBEC results.

**Figure 2.**
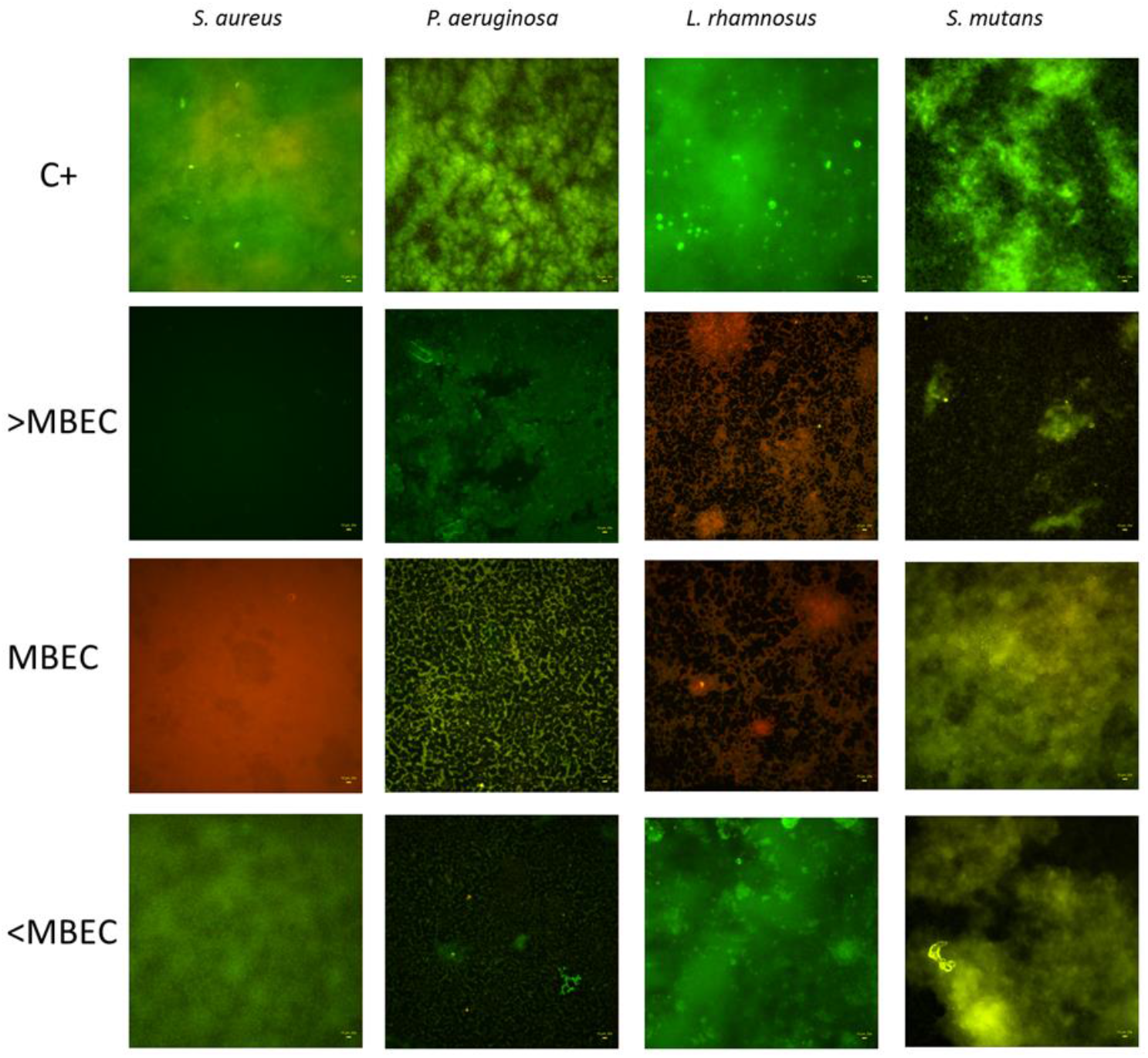
Fluorescent visualization of 4 tested biofilms after application MBEC, <MBEC and >MBEC concentrations of ASC1. Green staining represents live cells and red staining represents dead cells).

### 3.5. Biofilm Eradication from HAPd

Gram-negative strains displayed higher sensitivity to ASC1 than Gram-positive strains. In all cases, the eradication level exceeded 50% (**Figure 3A**). Simultaneously, various patterns of susceptibility to ASC1 were observed among tested *P. aeruginosa* strains (**Figure 3B**). Statistical significance was observed in the level of biofilm reduction between species from the same genera such as MRSA vs. *S. aureus* or *L. rhamnosus* vs. *L. casei* (**Figure 3C**).

**Figure 3.**
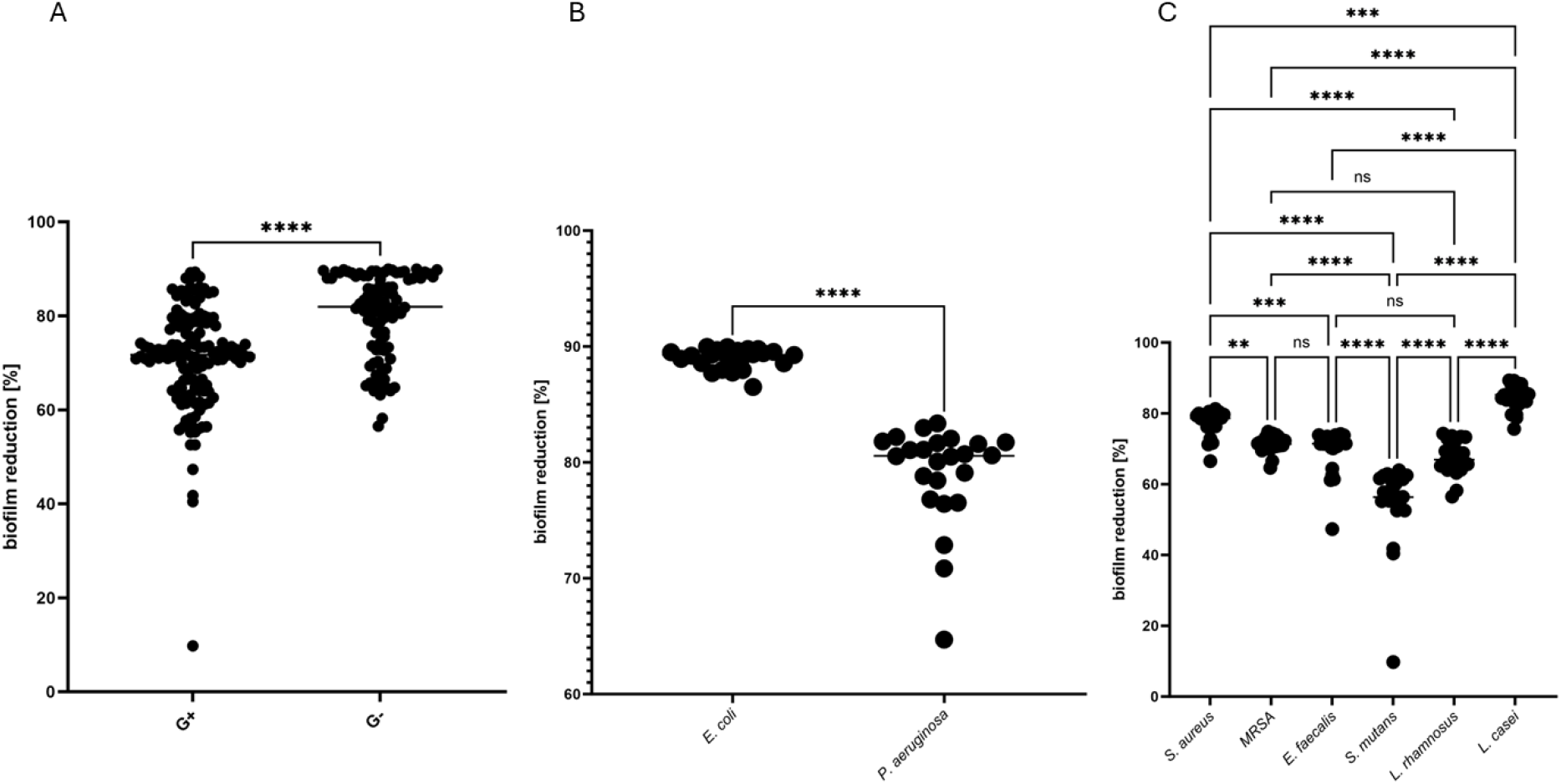
Eradication of biofilms formed by tested strains on HAPd.

ns - no statistical significance, **- small statistical significance, ***-moderate statistical significance, ****-strong statistical significance

**Figure 4.**
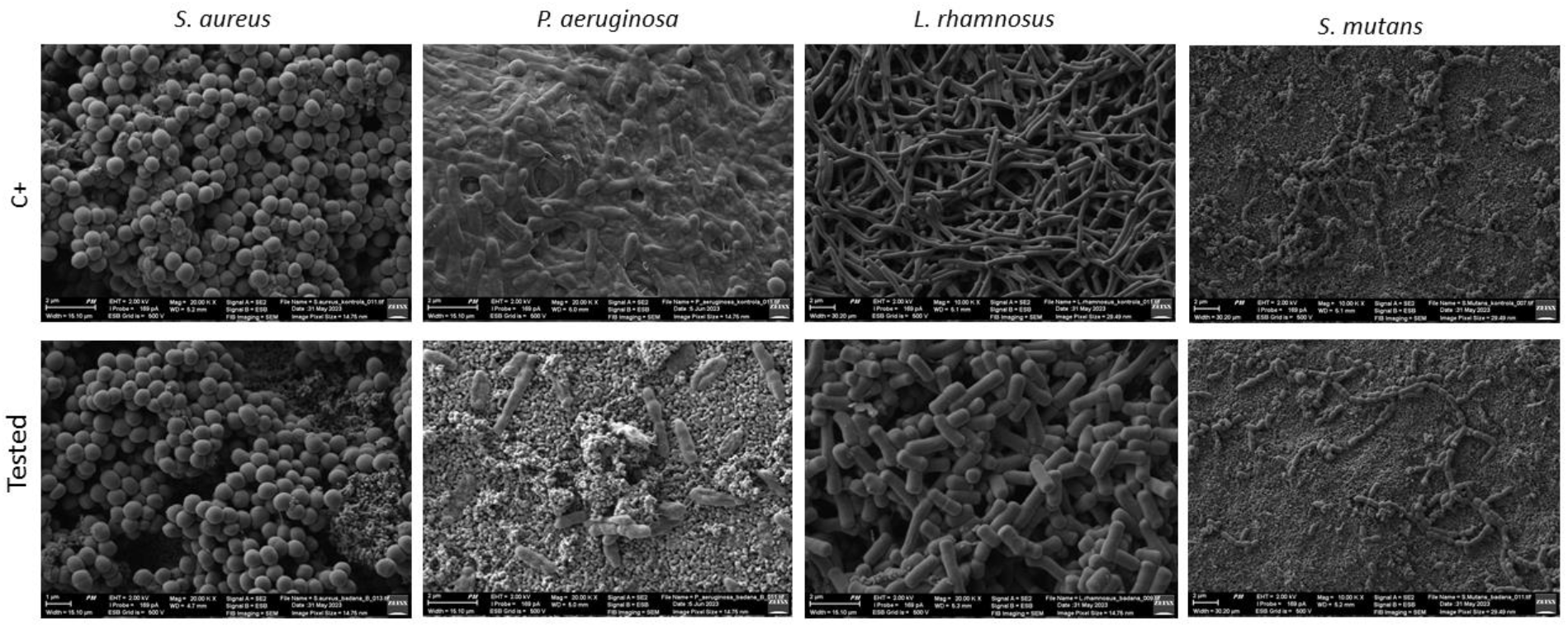
SEM visualization of biofilm formed on HAP surface before (C+) and after MBEC application. Magnification: *S. aureus* (x20 000), *P. aeruginosa* (x20 000), *L. rhamnosus* (x10 000), *S. mutans* (x10 000).

### 3.6. Visualization of biofilm with SEM

### 3.7. Evaluation of antibiofilm activity of ASC after different activation times

Obtained results (**Table 3**) revealed that activation time has an impact on the antibiofilm effectiveness of tested mixtures. Both ASC1 and ASC2 were characterized by greater antibiofilm activity after applying more than 60s of acidification time. However, extending acidification time over 180s did not translate into an increase in the antibiofilm activity of the tested solutions.

**Table 3.** MBEC values obtained after applying different times of activation of tested ASC.

|  | Acidification time [s] | MBEC MSSA |  | MBEC *P. aeruginosa* |  |
| --- | --- | --- | --- | --- | --- |
|  |  | %ClO<sub>2</sub> | ppm | %ClO<sub>2</sub> | ppm |
| **ASC 1** | 60 | 0.1915 | 1915 | 0.383 | 3830 |
|  | 180 | 0.047875 | 478.75 | 0.047875 | 478.75 |
|  | 360 | 0.047875 | 478.75 | 0.047875 | 478.75 |
| **ASC2** | 60 | 0.047875 | 478.75 | 0.383 | 3590 |
|  | 180 | 0.023938 | 239.38 | 0.047875 | 478.75 |
|  | 360 | 0.023938 | 239.38 | 0.047875 | 478.75 |

### 3.8. Cytotoxicity of Tested Solutions

#### A. in vitro

The cytotoxicity study showed that the solutions displayed similar cytotoxicity, regardless of whether SC was activated with an organic acid (GA) or an inorganic acid (HA). For both tested combinations, the lowest cytotoxic concentration was observed for ASC1 and ASC2 at 0.002992%, which demonstrated antimicrobial effectiveness against *S. aureus* and *E. coli* in MIC evaluation.

#### B. in vivo

The data on ASC1 cytotoxicity obtained in a two-dimensional model of fibroblast culture was developed using larvae *in vivo* model. Concentration A displayed rapid cytotoxic activity (exceeding the activity of highly concentrated ethanol), while concentration B did not lead to a significant drop in viability of larvae between hours 0 and 72 from the injection. After that time, c.a. 20% decrease in larvae viability was observed (**Figure 5**).

**Figure 5.**
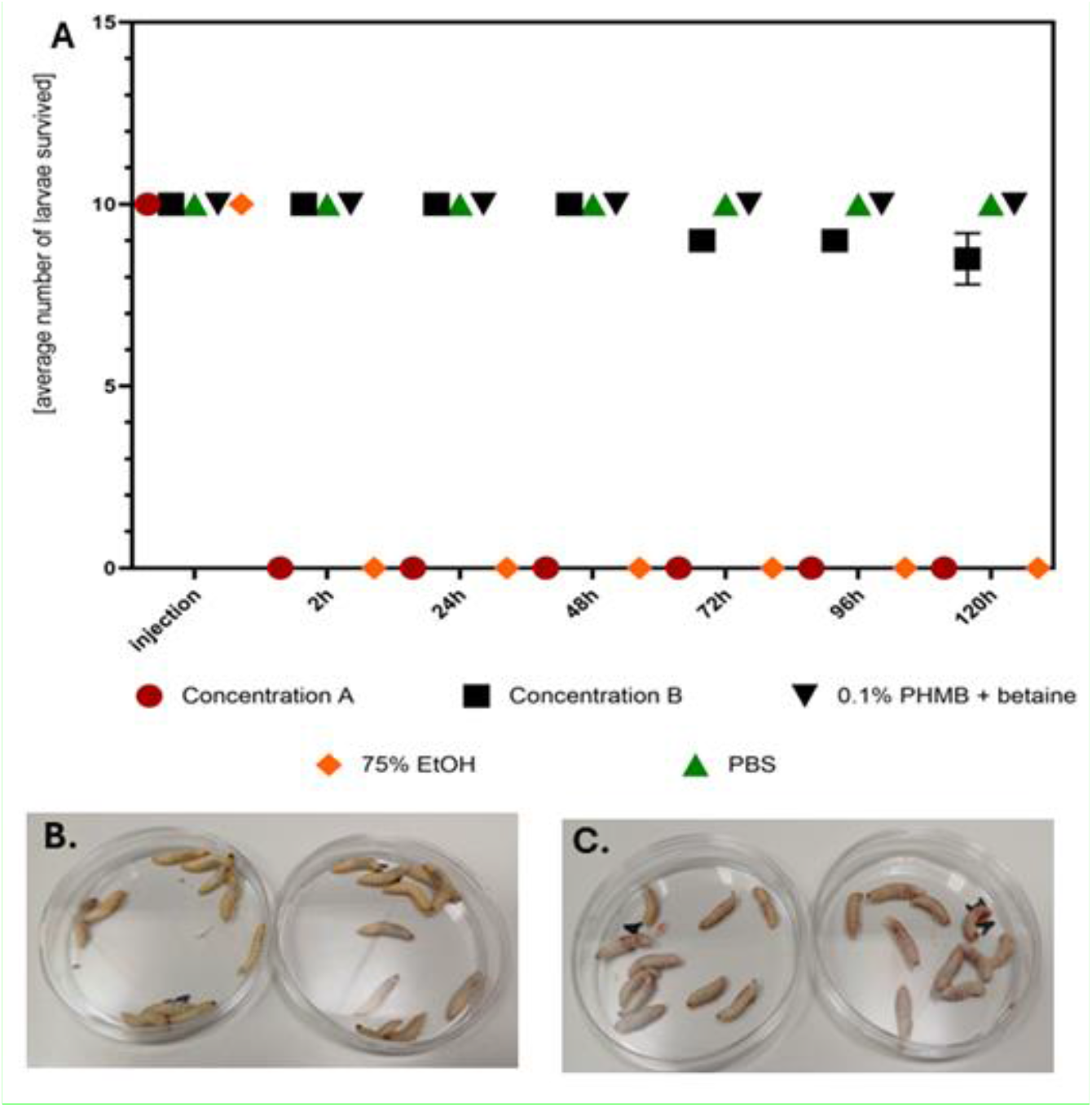
**A -** Average survival of larvae treated with concentration A (0.383% ClO_2_) or B (0.002992% ClO_2_) of ASC1, PHMB, PBS, and 70% ethanol **B -** larvae treated with concentration B of ASC1 display full turgor, mobility and viability. **C** – larvae treated with concentration A of ASC1 after two hours stopped moving and started to vomit red-colored liquid. Within the next hours these larvae became melanized and did not react to physical stimuli.

## 4. Discussion

ClO_2_ is known for its antimicrobial, antiviral, and antifungal activity when applied in controlled range of concentrations between 5 – 8000 ppm [1,22,23,29]. Numerous researchers explored its potential in *in vitro, in vivo* studies and clinical trials [17,23,24,29-32,33-36]. Also advocates of MMS assert its capability to treat a range of illnesses. However, such claims lack empirical support regarding both its safety and therapeutic effectiveness.

Herein, we aimed to investigate antimicrobial/antibiofilm activity and cytotoxicity of ASC, a source of ClO_2_ (**Table 1-4, Figure 1-4**). In our study, ClO2 was obtained by acidifying SC using either an inorganic or an organic acid (ASC1, ASC2, respectively). The lowest MIC of 0.002992% (29.92 ppm) was observed against MSSA and *E. coli* when ASC2 was applied. For ASC1, the same MIC value of 0.002992% (29.92 ppm) was observed only against MSSA (**Table 1**). Both formulations at this concentration showed acceptable levels of cytotoxicity but did not exhibit stronger antimicrobial activity than PHMB (the clinically used antiseptic agent) or the 5 and 20 ppm ASC solutions used by Ma et al. [29] (Table 4). The discrepancies observed may be related to either the different strains tested in this work or the ClO_2_ production method, as the electrolytic method reduces impurities in the solution, potentially affecting its antimicrobial efficacy [29, 35]. Conversely, MIC of PHMB was higher than of ASC2 towards *P. aeruginosa* and *L. casei* and of ASC1 towards *P. aeruginosa*. Our results are consistent with Morino et al. results which have shown the effectiveness of extremely low concentration (0.01 ppmv, 0.028 mg/m^3^) ClO_2_ against *P. aeruginosa* and *E. coli* [38]. In the case of *E. coli* and *L. casei*, ASC2 displayed a higher antimicrobial effect than ASC1. This may indicate that gluconic acid may have an additional role in the antibacterial effect [39,40]. The observed reduction in *L. casei* number may indicate that when drinking, ASC solution may disturb the gut microbiome and speculatively vaginal microbiota as well. All biofilms were more tolerant to applied compounds than their planktonic counterparts (**Table 1, 2**) [41,42]. *S. mutans* and *E. coli* biofilms were the most susceptible to ASC1 and ASC2. Accordingly, Herczegh et al. showed that 0.0015% and 0.003% solutions of ClO_2_ solutions are promising preventive and therapeutic adjuvants in dental practice [43]. SC was more effective against the biofilm of *S. mutans* than ASC1 and ASC2 (**Table 2, Figure 2**), which stays in line with reports showing that biofilm formation and matrix gene transcription can be stimulated by sublethal (4 μg/ml; 150ppm) doses of ClO_2_ [44,45].

**Table 4.**
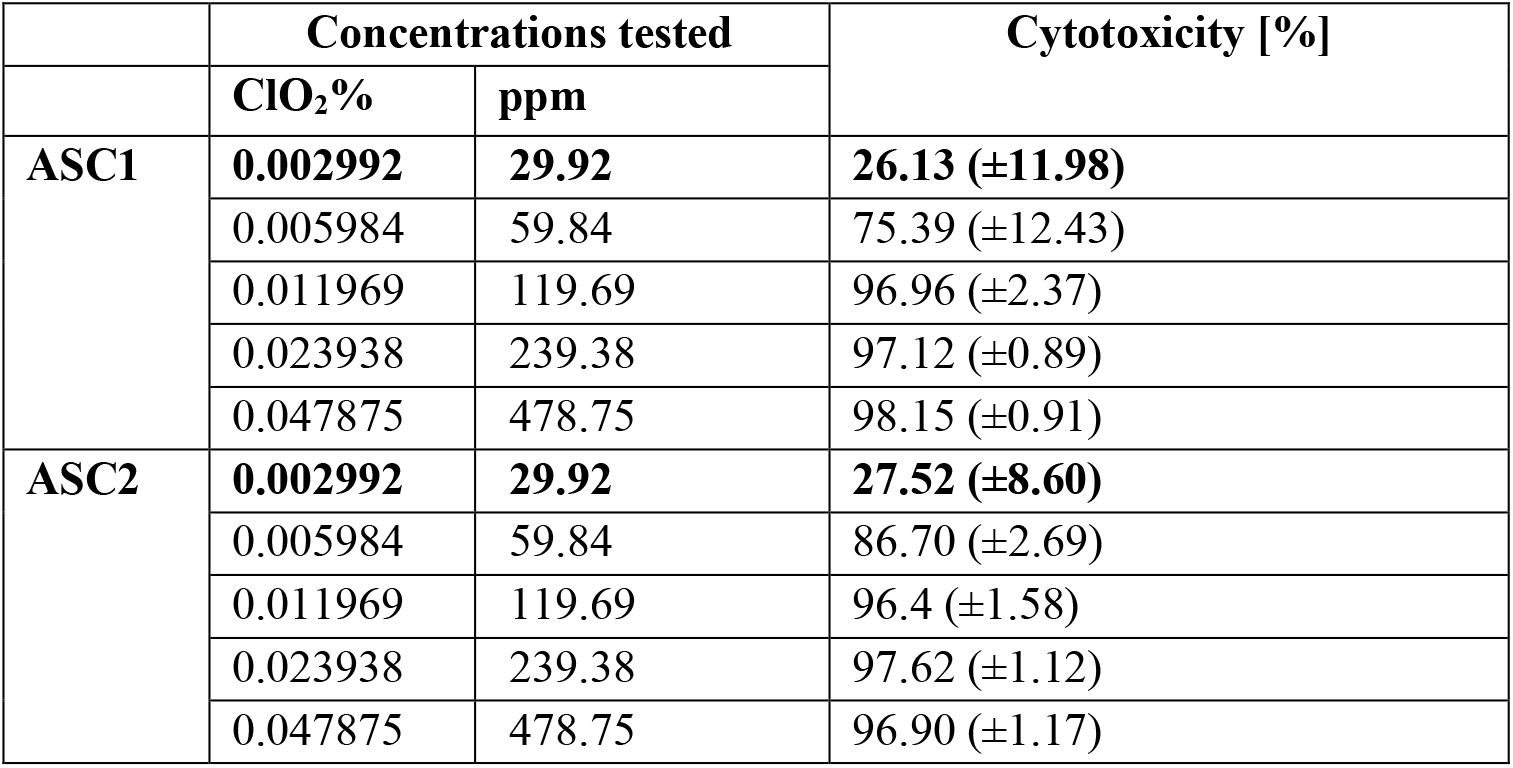
Cytotoxicity of selected concentrations of ASC1 and ASC2

It was shown that *L. rhamnosus* biofilm displayed high susceptibility to sub-MBEC (0.09575%, 957.5ppm) which may suggest that consuming even low amount of ClO_2_ may interfere negatively with microbiota (**Figure 2**), although contrary observations were also made by other authors [46].

The different species within *Lactobacillus* genera display various susceptibility to 75ppm and 125ppm ASC which has been explained by various adaptations to fermentative process conditions and so, different resistance to antimicrobial agents [47].

Gram-positive microorganisms demonstrated higher resilience to ClO_2_ than Gram-negative ones (**Figure 3**) which may be explained by the enhanced stability of the membrane structure and the mechanical stability of their cell walls [48,49,50].

It is believed that the duration of acidification of SC has an impact on its biological activity [3]. In this study, we shown that after 180s of acidification, effective concentrations of ASC were 2 - 8 times lower against MSSA and *P. aeruginosa* biofilms, respectively, compared to the situation when 60 s of acidification was applied. This result may influence the production of ASC solution for medical purposes [20-24]. To make ClO_2_, an acid is mixed with chlorite, which slowly releases the gas. The reaction normally requires high acidity (low pH) which is irritating for human tissues. Also in this study, we have observed a concentration-related increase of cytotoxic effects in both *in vitro* and *in vivo* analysis (**Table 4, Figure 5**) after the use of antimicrobially-active ASC concentrations. Choosing mild acid with a pH closer to neutral and prolonged acidification time may result in a product optimized for use on the body. Interestingly, the rapid death (within 2 hours) of larvae after application of ASC (**Figure 5**) was effect even harsher than level of cytotoxicity observed after exposure of fibroblast cell line to ASC.

## 5. Conclusions

Our results indicate that it is difficult to establish a precise antimicrobial dose of ASC while being effective against pathogens, safe for tissues and sparing probiotic species. We have shown that the dosage recommended by alternative medicine proponents cannot lead to the achievement of safe and therapeutic concentrations. At the same time, taking an ASC solution with a pH of 2.5 several times a day can led to damage to the mucous membrane of the gastrointestinal tract. Taking into consideration existing research results concerning ClO_2_ it can be noted that it has the potential to be used for medicinal purposes, but more investigation is needed to develop safe methods of its application and to evaluate a factual benefit/risk ratio.

## Transparency declarations

None to declare

